# Increasing absolute prey community density protects aposematic models and their imperfect Batesian mimics: Evidence from Neotropical *Adelpha* butterflies

**DOI:** 10.64898/2026.04.22.719628

**Authors:** Abigail E. Robinson, Alejandra Camargo-Cely, Julia B. Meyersiek, Colleen Fetherston, Sarah Speroff, Musfika Mishi, Kaela Sanborn, Maria Osipovich, Rachel Borzymowski, Jessica Herrmann, Susan Finkbeiner, Peter Buston, Sean P. Mullen

## Abstract

Batesian mimicry is a defensive adaptation where predators learn to avoid aposematic prey and generalize their warning signals to phenotypically similar mimics. The phenotypic accuracy needed for mimics to benefit from this adaptation depends on the relative densities of models and mimics and the model’s unpalatability. As aposematic models become more unpalatable or more common relative to their mimics, warning signals become stronger, allowing even poor mimics to benefit. However, few studies have disentangled the importance of *relative* frequencies of models and mimics from *absolute* density of the prey community (both models and mimics) in driving relaxed selection on imperfect mimics. Here, we test the hypothesis that increasing model unpalatability and absolute prey community density accelerates predator avoidance learning and enhances protection for imperfect mimics. Using replicas of the model *Adelpha iphiclus* (Linnaeus), its imperfect mimic *Adelpha serpa* (Boisduval), and the palatable control *Junonia evarete* (Cramer), we conducted field experiments that enhanced model unpalatability and doubled absolute prey density while maintaining a constant ratio of model, mimic, and control phenotypes. We found that enhanced model unpalatability and increased absolute density significantly reduced predation on all species, highlighting absolute community density as an underappreciated mechanism shaping selection on imperfect Batesian mimics.

## INTRODUCTION

Predator–prey interactions drive the evolution of diverse and complex anti-predator defenses and associated predator counterstrategies (Ruxton et al., 2018; Skelhorn et al., 2016; Speed & Turner, 1999). Within this broad coevolutionary framework, Batesian mimicry represents a classic predator avoidance strategy in which a palatable organism gains protection from its predators by resembling an unpalatable, or otherwise defended, model population (Bates, 1862). Although this classic understanding of Batesian mimicry predicts strong selection for close model–mimic resemblance, many Batesian mimics do not perfectly resemble their toxic models, raising important questions about the selective forces and ecological conditions that allow imperfect mimicry to persist (Penney et al., 2012; Ruxton et al., 2018; Sherratt, 2002). How perfectly a mimic must resemble its model depends on the foraging conditions of its predators (Sherratt & Peet-Paré, 2017). When predators have little incentive to discriminate between models and mimics, selection for precise resemblance is relaxed and predators generalize phenotypic traits more broadly (Edmunds, 2000; Kikuchi & Pfennig, 2013). Under this relaxed selection, mimics can deviate from their model’s appearance and still gain protection from predation, resulting in imperfect mimicry––where model–mimic resemblance is incomplete but functionally sufficient (Sherratt, 2011; Turner et al., 1984).

Theoretical, field, and laboratory studies show that the degree of predator generalization—and thus the level of phenotypic similarity needed to garner protection from predators—depends on factors such as the degree of model unpalatability, the relative abundances of models and mimics, and the availability of alternative prey (Brower et al., 1968; Kokko et al., 2003; Lindström et al., 2004; Pfennig et al., 2001; Ritland, 1994; Sherratt, 2002). Other works, both theoretical and empirical, highlight how environmental heterogeneity and predator and prey community complexity can also relax selection on imperfect mimics (Ihalainen et al., 2012; Kaczmarek et al., 2020; Kikuchi et al., 2022; Lindström et al., 1997, 2004; Luedeman et al., 1981; Malcolm, 1990; Sherratt & Peet-Paré, 2017). Despite these advances, key questions remain about the population- and community-level processes that allow imperfect mimics to persist in natural environments.

Aposematic prey––species that use bright colors or other conspicuous traits to warn predators about their toxicity––often vary in the strength of their chemical defense across natural environments (Brower et al., 1968; Ritland, 1995). This natural variation in chemical defense can correspond to changes in the dynamics of mimetic interactions (Prudic et al., 2019). Empirical studies show that mimics survive better when their models are highly unpalatable (Lindström et al., 1997; Pike & Burman, 2023), and that strongly defended models can accelerate predator avoidance learning (Skelhorn & Rowe, 2006). Theoretical work further demonstrates that increasing unpalatability strengthens the model’s warning signal and relaxes selection on mimetic accuracy, allowing imperfect mimics to persist despite copying the model’s phenotype imprecisely (Owen & Owen, 1984; Sherratt, 2002, 2011; Speed & Turner, 1999; Turner et al., 1984). In other words, when the cost of attacking a toxic model is sufficiently high, predators generalize warning signals more broadly, enabling imperfect mimics to remain sufficiently protected even without close resemblance to the model.

Just as greater model unpalatability enhances protection for imperfect mimics, increasing the abundance of a toxic model *relative* to its mimic also strengthens the model’s warning signal, thereby relaxing selection on high-fidelity mimicry (Edmunds, 2000; Kikuchi & Pfennig, 2013; Ruxton et al., 2018; Sherratt & Peet-Paré, 2017). When the toxic model is highly abundant relative to the mimic, predators face an increased risk of encountering it. Therefore, as the toxic model becomes more abundant relative to the mimic, predators experience stronger selection to avoid it, which also increases protection for mimics (Getty, 1985; Pfennig et al., 2001; Sherratt, 2011; Turner et al., 1984). Consistent with these theoretical predictions, several empirical studies show that mimics exhibit higher survival when their models have a higher relative abundance (Chouteau et al., 2016; Finkbeiner et al., 2018; Iserbyt et al., 2011; Lindström et al., 1997; Nelson et al., 2011; Pfennig et al., 2001). These effects of relative frequency are also visible in patterns of geographic variation in mimetic fidelity. For example, phenotypic resemblance between the eastern coral snake model and its Batesian mimic the scarlet kingsnake is higher near the model’s range edge, where the model is relatively rare, and lower in the center of the model’s range, where the model is relatively common (Harper & Pfennig, 2007). This pattern is consistent with the idea that selection for precise mimicry intensifies when the model is relatively rare, while poor mimics can persist when the model is relatively common because the strength of the model’s warning signal is heightened. On the other hand, when a palatable mimic is relatively more common than its toxic model, this can create a parasitic effect, where the model’s warning signal is so diluted by the high relative frequency of its mimic that survival of both models and mimics decreases (Aubier et al., 2017; Heerwig et al., 2023; Rowland, Mappes, et al., 2010; Skelhorn et al., 2011).

It is therefore well established that the relative frequency of toxic models and their palatable mimics plays a crucial role in determining when mimics are protected from predation. However, surprisingly little empirical work has disentangled these effects of *relative* frequency from those driven by *absolute* prey density. The role of absolute density in driving predator responses is well understood in the context of aposematic prey, where the strength of a warning signal increases with the rate at which predators encounter that signal, which increases as the absolute abundance of the aposematic organism increases (Chouteau et al., 2016; Endler & Mappes, 2004; Ioannou et al., 2008; Lotka & Volterra, 1925). As such, high absolute abundance reduces predation on defended aposematic prey (Alatalo & Mappes, 1996; Briolat et al., 2019; Finkbeiner et al., 2012; Kaczmarek et al., 2020; Kikuchi et al., 2021). Previous work has isolated relative model–mimic frequencies from total prey density, demonstrating that increasing the relative frequency of an unpalatable model, while holding total prey density constant, can enhance protection for imperfect Batesian mimics––thereby highlighting relative frequency as an independent mechanism driving mimic protection (Lindström et al., 1997). Yet it remains unclear whether changes in the absolute density of a mimetic community (models and mimics), with model–mimic ratios held constant, also affect the survival of aposematic and mimetic prey. Moreover, although several studies emphasize the importance of considering absolute density in experimental tests of mimicry, none have manipulated absolute prey density while maintaining constant model–mimic ratios (Lindström et al., 2001; Rowland et al., 2007; Rowland, Hoogesteger, et al., 2010). As a result, it is difficult to determine whether relaxed selection on imperfect mimics arises from shifts in relative frequency, changes in absolute community density, or a combination of the two.

Here, we investigate how model unpalatability and absolute prey density shape protection for imperfect Batesian mimics in Neotropical *Adelpha* butterflies, a phylogenetically diverse genus characterized by extensive convergent evolution of wing patterns and the formation of multi-species mimicry rings composed of both palatable and unpalatable taxa (Ebel et al., 2015; Mullen et al., 2011; Páez V. et al., 2025; Willmott, 2003). In *Adelpha*, unpalatable models typically specialize on toxic *Rubiaceae* host plants and likely sequester defensive secondary metabolites from these plants, whereas palatable mimics are generalists that feed on non-*Rubiaceae* hosts (Finkbeiner et al., 2017; Kessler & Baldwin, 2002; Lopes et al., 2004; Prudic et al., 2007; Soto-Sobenis et al., 2001).

A well-studied pair within this system is the toxic *Rubiaceae* specialist *Adelpha iphiclus* (Linnaeus) and its near-identical Batesian mimic *Adelpha serpa* (Boisduval) (Finkbeiner et al., 2017, 2018). Despite their striking phenotypic similarity, field experiments in Costa Rica showed that while predators avoided *A. iphiclus*, *A. serpa* was not protected from predation, indicating that predators were able to discriminate between the two and selectively forage only on the edible mimic (Finkbeiner et al., 2017). This predator discrimination, despite near-identical wing patterns, renders *A. serpa* an imperfect mimic. Subsequent work demonstrated that protection for this imperfect mimic is governed by negative frequency dependent selection: while *A. serpa* is not protected in Costa Rica, where the species occurs at a similar frequency to its model, *A. serpa* is protected in Ecuador, where it is much rarer relative to its model (Finkbeiner et al., 2018). This work highlights the central role of relative model–mimic frequencies in maintaining mimicry in *Adelpha* and positions *Adelpha* as a powerful system for testing how absolute prey density and model toxicity interact to shape selection on mimicry complexes.

Using field predation experiments in Costa Rica, we test how model unpalatability and absolute prey density impact survival of the chemically defended model *Adelpha iphiclus*, its palatable, Batesian mimic *Adelpha serpa*, and a palatable control *Junonia evarete* (Cramer). Experiments were carried out using artificial butterfly facsimiles at the same Costa Rican field site used by Finkbeiner et al. (2017). To isolate absolute density effects from relative frequency effects, we doubled the absolute facsimile prey density while maintaining a constant 1:1:1 ratio of model (*A. iphiclus*), mimic (*A. serpa*), and control (*J. evarete*) facsimiles. We hypothesized that, at a fixed model–mimic ratio, increasing absolute prey density and model unpalatability would accelerate predator avoidance learning (Hypothesis 1) and increase protection for the imperfect mimic (Hypothesis 2). To test this, we designed an experiment with two levels of absolute prey density (low vs. high) and two levels of model unpalatability (unaltered vs. experimentally enhanced), enabling us to investigate important questions about how model unpalatability and absolute prey density interact to relax selection on mimetic communities.

## METHODS

### Field site

To explore how changes in model unpalatability and absolute prey density impact protection for mimetic butterflies, we conducted a field predation experiment at the Organization for Tropical Studies’ (OTS) La Selva Biological Reserve (10°25’28”N, −84°0’18”W) in Sarapiquí, Costa Rica between May and June 2022. La Selva contains 1,600 hectares of old growth and secondary growth tropical lowland forest bordering the Northern section of Braulio Carrillo National Park. As such, this reserve offers unparalleled access to a diverse and abundant community of insectivorous avian predators. Several large-scale field experiments using similar artificial butterfly replicas have been completed at La Selva, demonstrating that this field site is suitable for conducting predation experiments of this nature (Finkbeiner et al., 2012, 2017, 2018).

### Butterfly facsimile production

Artificial butterfly facsimiles were used to measure predation across experimental treatments. We used color-accurate facsimiles of *Adelpha iphiclus* (model), *Adelpha serpa* (mimic), and *Junonia evarete* (control) designed by Finkbeiner et al. (2017). In Finkbeiner et al. (2017), spectral reflectance measurements were taken and compared against real butterfly wings for each species, revealing no differences in discriminability between facsimiles and real butterfly wings. These facsimiles were successfully used to assess the effects of relative frequency on predator learning within the same *Adelpha* mimicry system in the same field site (La Selva). Therefore, no changes were made to facsimile design in the present study, and additional spectral reflectance measurements were not taken.

Facsimiles were constructed following protocols outlined in Robinson et al. (2025). Facsimile images were double-printed using an Epson Stylus Pro 4900 printer with UltraChrome® High Dynamic Range ink on Grade 1 Whatman filter paper (Finkbeiner et al., 2012, 2017; Robinson et al., 2025). Once printed, facsimiles were pasted onto a cardstock backing (brand: Bazzil®; color: mocha divine) using Krylon® high-strength spray adhesive. After 48 hours, facsimiles were cut out and sprayed with Krylon® Matte Finish, which provided moisture resistance and protection from color damage. Black twist ties were attached to each facsimile, which allowed them to be easily affixed to foliage and branches in the field. Finally, abdomens were constructed out of black Newplast® plasticine and attached to each facsimile. Plasticine abdomens are often used in field predation experiments, and have been shown to allow for effective detection of predation events through impressions left in the clay when predators attempt to bite the facsimiles (Kikuchi & Pfennig, 2010; Niskanen & Mappes, 2005).

### Predation experimental design

Using paired absolute prey density (low vs. high) and model unpalatability (unaltered vs. experimentally enhanced) treatments across 100 independent field sites at La Selva Biological Reserve, we assessed how variation in absolute prey density and palatability of the chemically defended model *A. iphiclus* impacts predation rates on all facsimile types (Figure 1). Three different butterfly facsimiles were included in all treatments: (1) *Adelpha iphiclus*, the toxic model, (2) *Adelpha serpa*, the palatable Batesian mimic, and (3) *Junonia evarete*, a common palatable butterfly species that was used as a control. To isolate the effect of absolute prey density on facsimile survival, we maintained a 1:1:1 ratio of *A. iphiclus*, *A. serpa,* and *J. evarete* phenotypes in all field sites, and doubled absolute density of all prey types in high-density treatments relative to the low-density treatments (Figure 1). For low-density treatments, 20 facsimiles of each butterfly species were placed in each field site (*i.e.,* 60 facsimiles total/site). Total facsimile density was doubled in high-density treatments, with 40 facsimiles of each species in each field site (*i.e.,* 120 facsimiles total/site).

**Figure 1.**
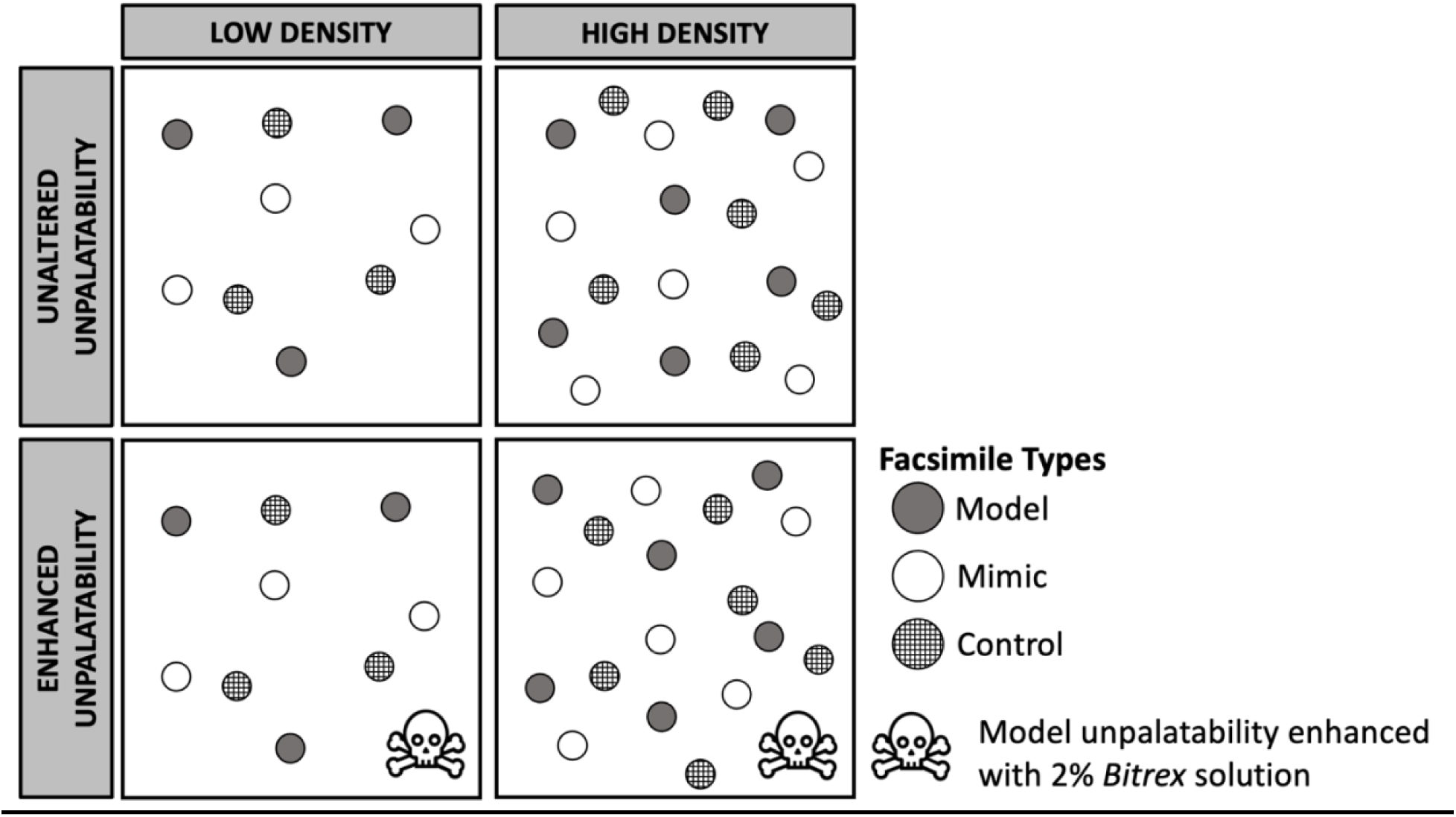
Schematic representation of paired density (low vs. high) and unpalatability (unaltered vs. enhanced) experimental treatments. To isolate absolute density effects on predator foraging, the relative frequency of model and mimic and control facsimiles was held constant at a 1:1:1 ratio across treatments and absolute prey density was doubled in the high-density treatments. **Alt Text:** Graphical representation of paired experimental design with unaltered vs. enhanced unpalatability treatments and low vs. high density treatments where absolute community density is doubled in high density treatments

Low- and high-density treatments were paired with unaltered and experimentally enhanced model unpalatability treatments to evaluate how model unpalatability mediates the effect of prey density on predator foraging decisions (Figure 1). In the enhanced unpalatability treatments, facsimiles of *A. iphiclus* were sprayed with a 2% Bitrex solution (denatonium benzoate) to enhance this species’ naturally occurring chemical defense. Bitrex is an extremely bitter compound that has been shown to effectively drive avoidance learning in avian predators (Curley et al., 2015; McLellan et al., 2023; Siddall & Marples, 2008; Winsor et al., 2020). Bitrex was reapplied to *A. iphiclus* facsimiles daily in all enhanced unpalatability field sites. Palatability was not experimentally manipulated in treatments with unaltered model unpalatability.

Each treatment (low-density, no Bitrex; low-density, Bitrex; high-density, no Bitrex; and high-density, Bitrex) was replicated in 25 independent field sites at La Selva Biological Reserve, resulting in 100 total field sites used in this experiment. Therefore, each low-density treatment involved 500 facsimiles per species and 1,500 facsimiles total, while each high-density treatment involved 1,000 facsimiles per species and 3,000 facsimiles total. Field sites were separated by at least 250 meters, which aligns with estimated avian home range sizes in the Neotropics (Samuel et al., 1985; Wang et al., 2007). This design minimizes the possibility that a small group of predators were responsible for all facsimile attacks, and enables us to sample across independent predator communities (Finkbeiner et al., 2012; Kristiansen et al., 2018). Within each site, facsimiles were attached to foliage at least 1 meter high with dorsal wings facing upward. This placement resembles perching behavior and ensures that facsimiles were visible to avian predators. All facsimiles were checked daily for evidence of avian predation, which is often characterized by triangular beak-shaped or deep puncture marks (Figure 2). Suspected predation events were photographed, and facsimiles were reset daily by smoothing or replacing plasticine abdomens. A maximum of one predation event was recorded for each facsimile on each day of our experiment, and all facsimiles were removed after four days. All photographs were evaluated by a team of researchers and categorized independently as “avian predation” or “non-avian predation”.

**Figure 2.**
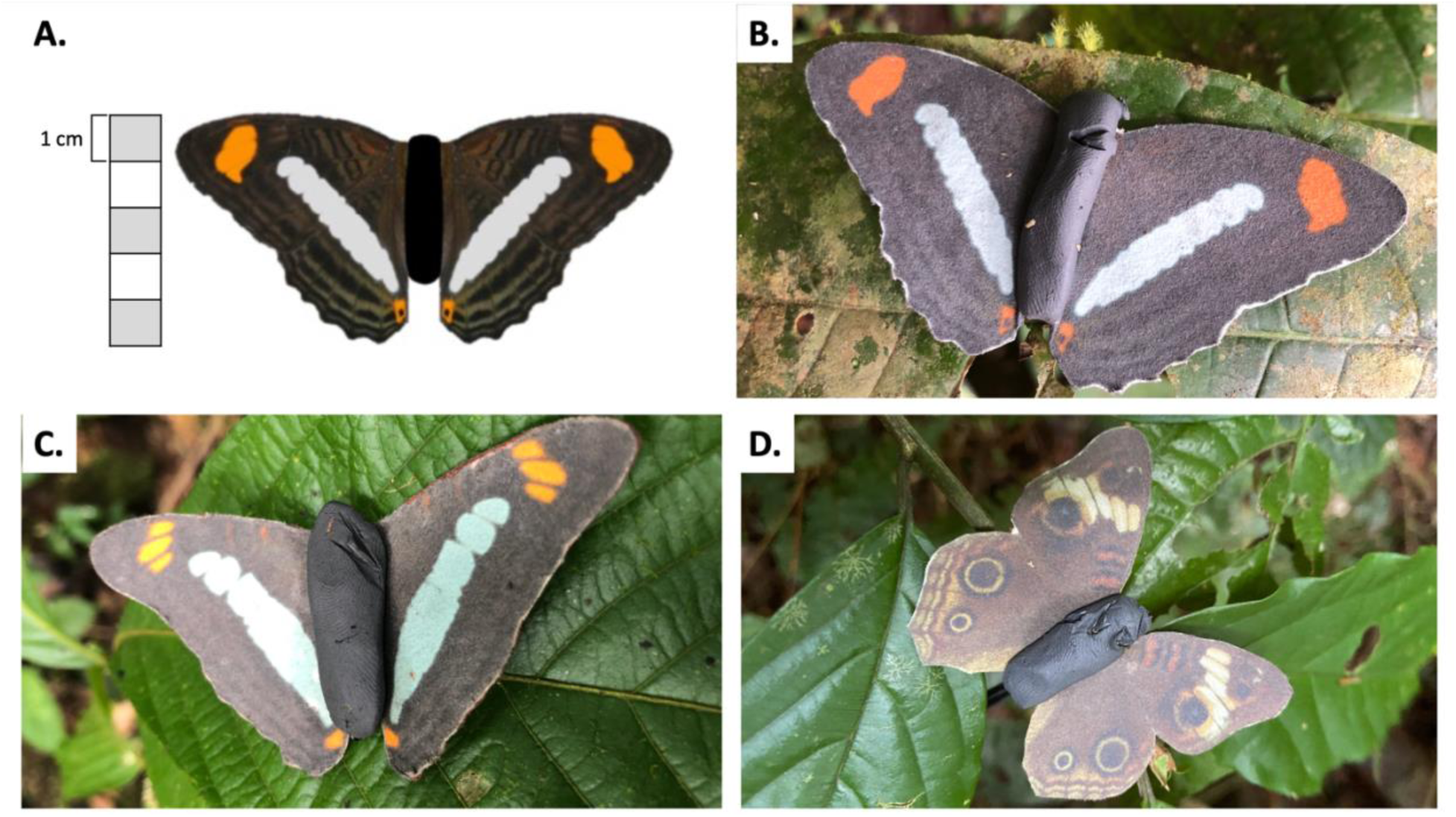
(A) Approximate size of *A. iphiclus* facsimiles. Wingspan was consistent across facsimile types. Images show evidence of avian predation on facsimiles in the field for *A. iphiclus* (B), *A. serpa* (C), and *J. evarete* (D). **Alt Text:** Panels A through D show images of facsimiles. Panel A illustrates that facsimiles are approximately 5 cm in height, while panels B through D show images of *A. phiclus*, *A. serpa*, and *J. evarete* facsimiles, respectively. All images show evidence of avian predation, which is identified through triangular beak-shaped marks or deep puncture marks in clay absomens.

### Statistical analysis

To test the hypothesis that increased absolute prey density and enhanced model unpalatability accelerate predator avoidance learning (H1), we fitted a mixed-effects Cox proportional hazards model using the *survival* and *coxme* packages (T. Therneau, 2015; T. M. Therneau, 2001). This time-to-event modeling approach allows the rate at which predation events occur to vary across levels of a response variable. Sites that experienced no predation events during the four-day experiment were excluded from all analyses. The rationale for this exclusion is that the absence of attacks across all facsimile types at these sites likely reflects a lack of insectivorous predator activity rather than treatment-specific behavioral responses, and therefore their inclusion could bias estimates of attack rates. Our initial model included species, absolute density, model unpalatability, and all two-way and three-way interactions as predictor variables to test whether the interactive effects of density and unpalatability differed across species. Following model selection, the final model included only the fixed effects of species, absolute density, and model unpalatability, with field site included as a random effect.

To test the hypothesis that increased absolute prey density and enhanced model unpalatability reduce overall predation on aposematic models and their Batesian mimics (H2), we fitted a generalized linear mixed-effects model (GLMM) with a binomial distribution using the *glmmTMB* package (Brooks et al., 2017). This analysis used the same subset of data as the survival analysis (*i.e.,* excluding sites with no recorded predator activity). Following the same model selection process outlined for the Cox proportional hazards model, the final GLMM included species, absolute density, and model unpalatability as fixed effects, and field site as a random effect.

Model selection for both analyses was based on comparisons of Akaike’s Information Criterion (AIC) values and likelihood ratio tests conducted using the *anova* function. Model comparisons are presented in Table S1 (Cox proportional hazards model) and Table S2 (GLMM). Model assumptions for the GLMM were evaluated using the *DHARMa* package (Hartig, 2022), which indicated no significant deviations from expectations under a binomial distribution. For both models, we conducted pairwise contrasts of species using the *emmeans* package (Lenth, 2023), and applied a Holm p-value correction to account for multiple comparisons. All analyses for both models were performed with R statistical software (R Core Team, 2022).

## RESULTS

Across 100 independent field sites, 86 experienced at least one avian predation event (25 in the high-density, no Bitrex treatment; 22 in the high-density, Bitrex treatment; 19 in the low-density, no Bitrex treatment; and 20 in the low-density, Bitrex treatment). All subsequent analyses are based on these 86 field sites, with 14 field sites excluded.

Considering our first hypothesis regarding the rate of learning, we found that absolute density, model unpalatability, and species all affected the rate at which predators learned to avoid warning signals (Figure 3; Table 1). Specifically, our mixed-effects Cox proportional hazards model showed that avian predators learn to avoid facsimiles significantly faster under high absolute density conditions (χ^2^ = 5.5503, p = 0.01848) and under enhanced model unpalatability conditions (χ^2^ = 14.1156, p = 0.00017). We also found a significant main effect of species (χ^2^ = 9.8344, p = 0.00732), and our analysis of pairwise contrasts demonstrated that predators learn to avoid *A. iphiclus* (model) significantly faster than *A. serpa* (mimic; p = 0.0193) and *J. evarete* (control; p = 0.0090).

**Figure 3.**
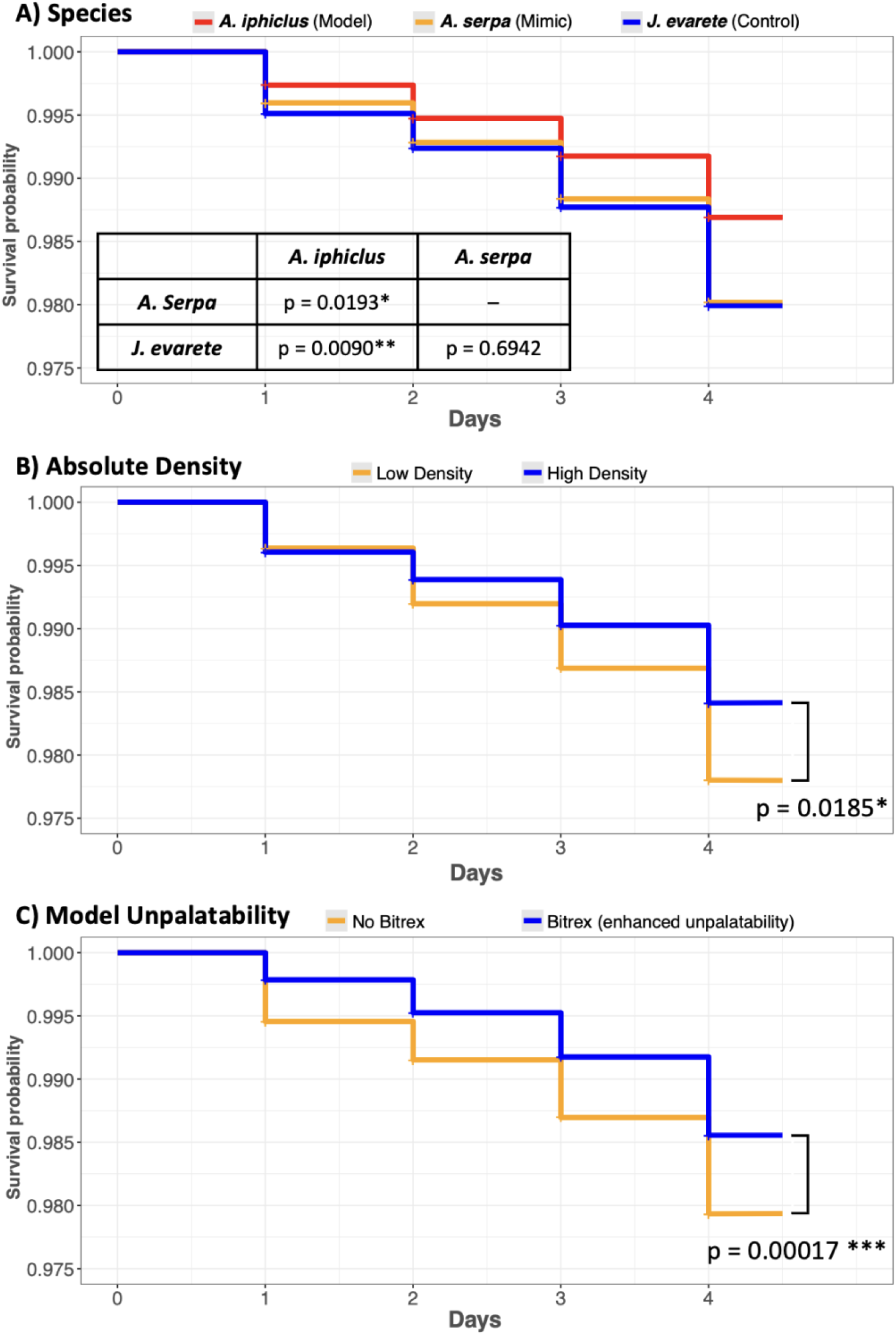
Kaplan-Meier survival curves comparing the fixed effect of (A) species, (B) absolute density, and (C) model unpalatability on facsimile survival, demonstrating the speed of predator avoidance learning across predictor variables. Asterisk indicates significant difference (p < 0.05) in survival probabilities based on mixed-effects Cox proportional hazards model and pairwise comparisons across species. **Alt Text**: Three-panel graph showing the survival probability of facsimiles over time across treatments with significance values. For all panels, the x-axis shows days, and the y-axis shows survival probability. Panel A shows that *A. iphiclus* had a higher survival probability than both *A. serpa* and *J. evarete*. Panel B shows that predators learned to avoid facsimiles significantly faster under high density conditions compared to low density conditions, and panel C shows that predators also learned to avoid facsimiles significantly faster under conditions of high model unpalatability relative to treatments with untreated model facsimiles.

**Table 1.**
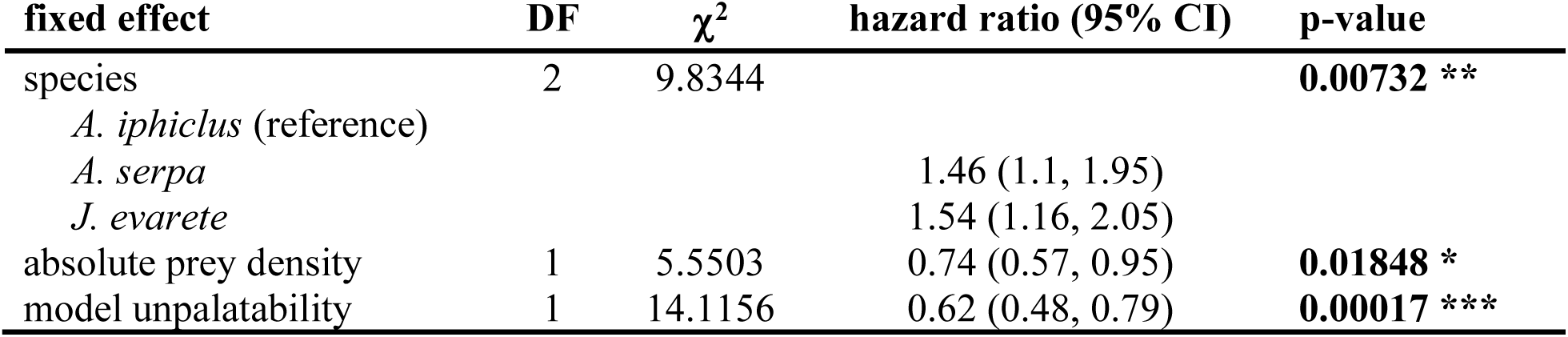
Results of best fit mixed-effects Cox proportional hazards regression model evaluating effect of species, absolute prey density, and model unpalatability on facsimile attack rates over time as a proxy for predator learning speed. Significant p-values are shown in bold (* p < 0.05, ** p <0.01, *** p < 0.001)

Considering our second hypothesis regarding overall predation rate, we found that absolute density, model unpalatability, and species all affected the overall predation rate. Specifically, our generalized linear mixed effect model revealed significant main effects of absolute density (χ^2^ = 5.4996, p = 0.0190212), model unpalatability (χ^2^ = 14.1333, p = 0.0001703), and species (χ^2^ = 9.9402, p = 0.0069426) on overall predation rates (Figure 4; Table 2). We found that all facsimile types were attacked significantly less under high absolute density conditions and conditions of enhanced model unpalatability (Figure 4). We also found that facsimiles of *A. iphiclus* experience significantly less predation than facsimiles of *A. serpa* and *J. evarete* (Table 2).

**Figure 4.**
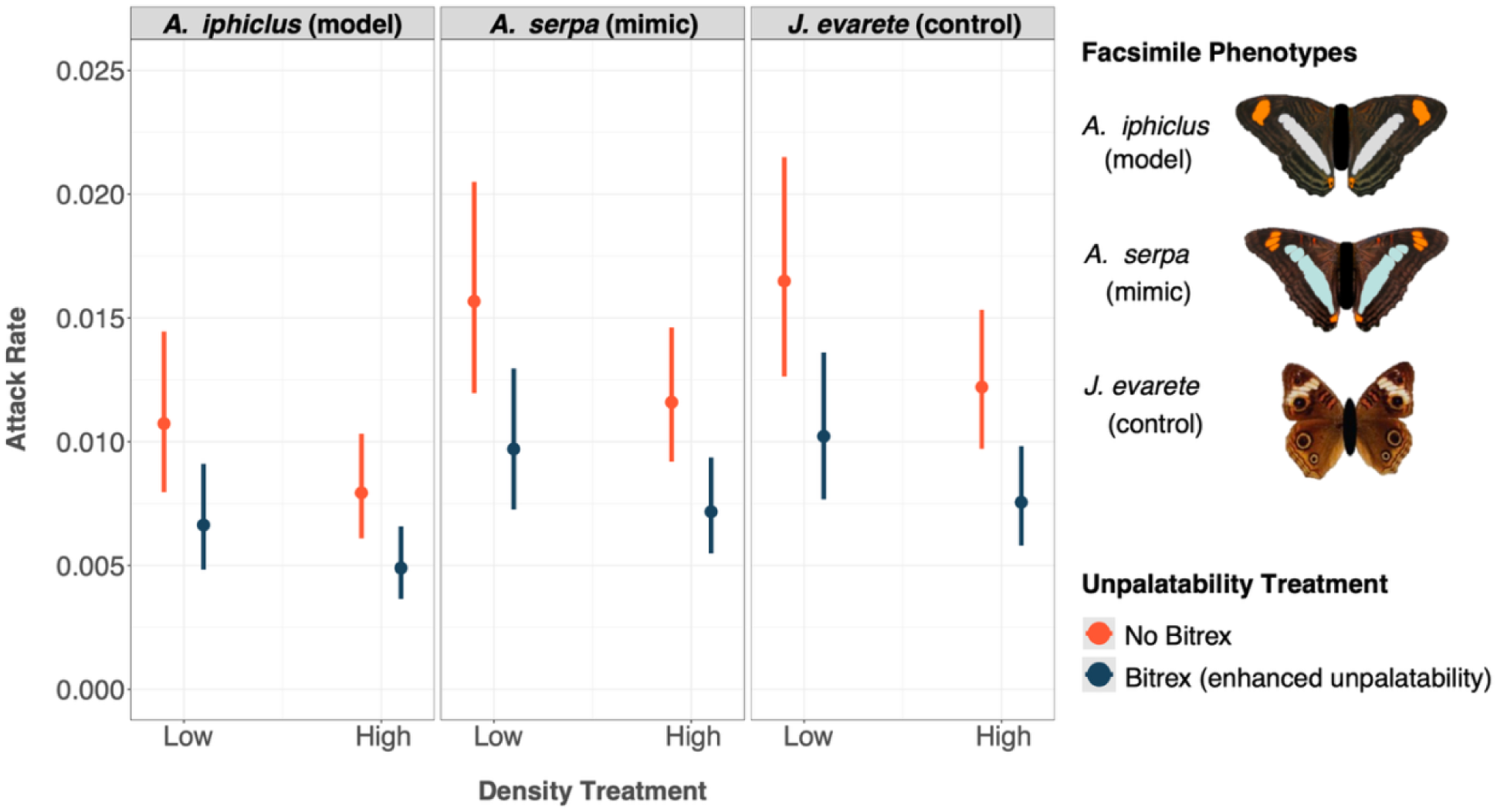
Estimated fixed effects of species, density, and unpalatability on facsimile attack rates based on results of generalized linear mixed model with binomial distribution. Color indicates unpalatability condition (orange = no added unpalatability, blue = unpalatability of *A. iphiclus* facsimiles was enhanced with a 2% Bitrex solution). Error bars represent 95% confidence intervals. **Alt Text**: Three-panel dot and whisker graph showing overall attack rates on each species; the x-axis shows density treatment (low vs. high) and the y-axis showed attack rate across all days. Dot and whisker plots are colored by model unpalatability treatment (no Bitrex vs. Bitrex).

**Table 2.**
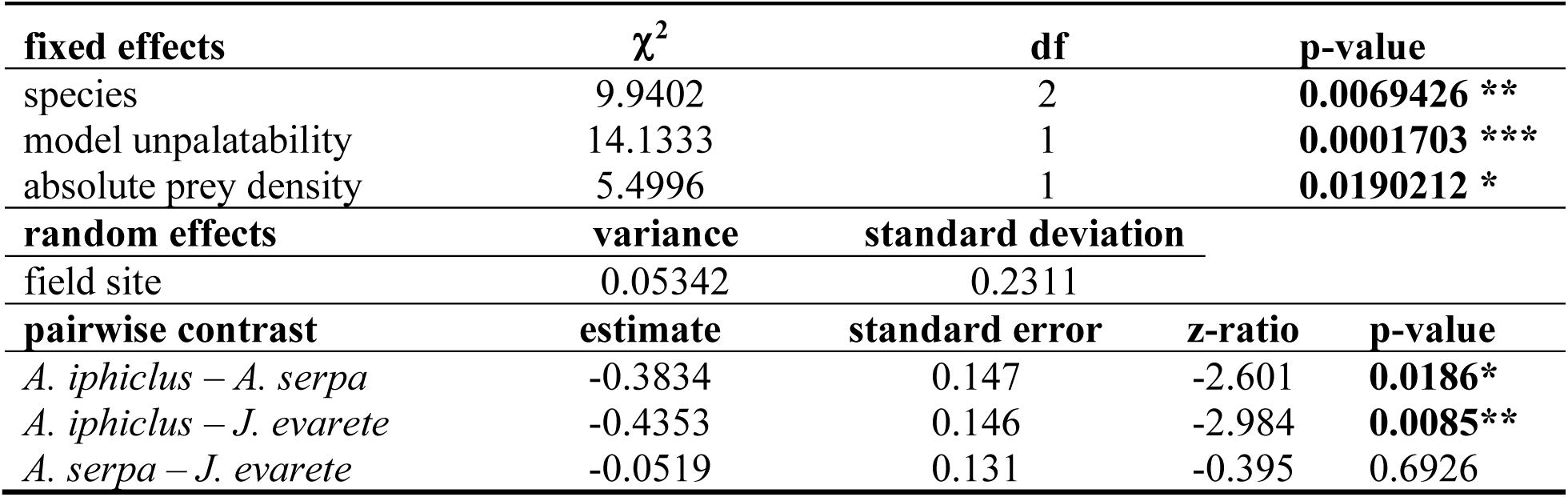
Results of best fit generalized linear mixed model with binomial distribution testing fixed effect of species, model unpalatability, and absolute prey density on facsimile attack rates. Field site was included as a random effect. Results of pairwise contrasts of species are shown; p-values for all pairwise comparisons were adjusted using the Holm method for multiple comparisons. Significant p-values are shown in bold (* p < 0.05, ** p <0.01, *** p < 0.001)

## DISCUSSION

We hypothesized that increasing absolute prey density and enhancing model unpalatability would i) accelerate predator avoidance learning and ii) reduce overall predation pressure on the *Adelpha* mimicry complex. Consistent with this prediction, higher absolute prey density, while holding model–mimic relative frequencies constant, increased the rate at which predators learned to avoid our facsimiles and reduced overall predation on the facsimile prey community. Similarly, enhanced model unpalatability accelerated predator learning and lowered predation rates. The combination of high absolute density and enhanced model unpalatability produced the lowest overall predation on the mimicry complex, suggesting that greater absolute density together with stronger chemical defense most effectively amplified the model’s warning signal.

Across all treatments, predators learned to avoid *A. iphiclus* (model) faster than *A. serpa* (mimic) and *J. evarete* (control), and the defended model experienced the lowest predation overall. However, we did not find evidence of a significant interaction between species, absolute density, and model unpalatability, suggesting that all three species (model, mimic, and control) benefitted equally from these enhanced warning signals. These findings align with previous work showing that greater model unpalatability can accelerate predator learning and relax selection on imperfect mimics (Ihalainen et al., 2007; Lindström et al., 1997; Pike & Burman, 2023). The comparable benefit observed for the palatable control further suggests that the protective effects of heightened warning signals may extend to particularly poor mimics, which aligns with previous work demonstrating that protection for poor mimics is greater in complex environments with high signal diversity (Beatty et al., 2004; Ihalainen et al., 2012; Sherratt & Peet-Paré, 2017).

Although treatment effects did not differ significantly among species, predators consistently learned to avoid the chemically defended model more quickly than either the mimic or the control, and the model maintained the lowest predation rates across all treatments. Thus, while strengthening the model’s warning signal enhanced protection for imperfect mimics, aposematic prey still derived the greatest benefit. This pattern is consistent with the expectation that stronger warning signals elicit stronger avoidance responses (Skowron Volponi et al., 2025), and previous work demonstrating that increasing absolute density increases protection for aposematic prey (Briolat et al., 2019; Jeschke & Tollrian, 2000). However, our results do not align with studies suggesting that Batesian mimicry can be parasitic (Aubier et al., 2017; Heerwig et al., 2023). In contrast, we found that increasing absolute prey density while maintaining constant relative frequencies reduced predation on both the aposematic model and its imperfect mimic. This result suggests that mimicry may only become parasitic when mimics are relatively more abundant than their models and that increasing the absolute density of the mimicry complex may permit increased mimetic abundance without generating parasitic effects (Rowland, Mappes, et al., 2010). To our knowledge, this study represents the first field-based predation experiment to isolate absolute density effects on predator foraging by increasing absolute prey density while maintaining a constant 1:1:1 ratio of models, mimics, and controls.

Negative frequency dependent selection shapes protection for imperfect mimics, where decreasing the abundance of a mimic relative to its defended model increases protection for the mimic (Pfennig et al., 2001; Ruxton et al., 2018). This pattern has been documented in several mimicry systems, including the *Adelpha* mimicry complex (Chouteau et al., 2016; Finkbeiner et al., 2018; Shine et al., 2022). Previous work has shown that in Costa Rica, avian predators can distinguish between the toxic model *A. iphiclus* and the imperfect mimic *A. serpa* despite their striking phenotypic similarity, likely because both species occur at similar population densities (Finkbeiner et al., 2017, 2018). In contrast, protection for *A. serpa* increases in Ecuador, where this Batesian mimic is much rarer relative to its model, underscoring the role of relative abundance in driving relaxed selection on imperfect mimics (Finkbeiner et al., 2018). Negative frequency dependent selection is thought to operate by increasing the rate at which predators encounter the model relative to the mimic, thereby strengthening the model’s warning signal (Ioannou et al., 2008; Lindström et al., 1997; Luedeman et al., 1981). Our results suggest that increasing the absolute density of a mimetic community, and thereby increasing predator encounter rates with all prey types equally (Endler & Mappes, 2004; Ioannou et al., 2008; Mols et al., 2004), can similarly strengthen the model’s warning signal. This finding indicates that a higher absolute abundance of mimics may not dilute the model’s warning signal when relative frequencies are maintained. As such, we highlight an underappreciated role for the absolute density of a mimicry complex, independent of model–mimic relative frequencies, in shaping relaxed selection on imperfect mimicry.

Although our experiment did not directly account for alternative prey availability or predator hunger, both factors likely influenced predator decision-making and represent important limitations of this study. When alternative prey are abundant, the immediate payoff of sampling imperfect mimics is low, reducing the incentive for predators to invest in learning fine-scale distinctions between aposematic models and their mimics and favoring broad generalization of warning signals (Kokko et al., 2003; Lindström et al., 2004; Sherratt & Peet-Paré, 2017). La Selva supports a highly diverse insect community that likely provides ample alternative food sources for insectivorous birds (MacDade et al., 1994). While we did not directly measure the density of alternative prey at La Selva, this ecological context suggests that predators in our field sites could afford to generalize avoidance of the model (Dill, 1975; Sherratt, 2003), driving reduced predation on *A. iphiclus,* its mimic *A. serpa*, and the palatable control *J. evarete* in our experiment when that model’s warning signal was strengthened. Together, these patterns are consistent with an information-acquisition mechanism underlying relaxed selection on imperfect Batesian mimics. Although avian predators are capable of fine-scale discrimination between similar phenotypes—including *A. iphiclus* and *A. serpa* (Finkbeiner et al., 2017; Langham, 2004; Llaurens et al., 2014) they are expected to use this information only when the benefits outweigh the costs (Sherratt & Peet-Paré, 2017). The observed reduced predation on the palatable control when the model’s warning signal was strengthened suggests that increasing the costs associated sampling facsimiles can promote global avoidance, with predators rejecting all facsimiles in favor of alternative food sources.

Predator motivation to distinguish between models and mimics is also likely influenced by hunger levels and the relative nutritional value of mimetic prey compared to alternative food sources (Halpin et al., 2014; Kaczmarek et al., 2020; Kokko et al., 2003; Luedeman et al., 1981). Predators have been shown to forage on toxic prey or learn to differentiate toxic prey from phenotypically similar mimics when hunger is extreme, when defended prey offer high nutritional rewards, or when alternative food sources are scarce (Barnett et al., 2007; Halpin et al., 2014; Kokko et al., 2003; Lindström et al., 2004; Skelhorn & Rowe, 2007; Veselý et al., 2016). While our findings provide novel insight into absolute density as a mechanism underlying imperfect mimicry, our experimental design did not allow us to assess how variation in predator hunger or prey nutritional value modulates these dynamics. Our experiment did not provide predators with a food reward, which may have contributed to the extreme generalization we observed. Indeed, if our facsimiles had provided nutritional value to predators, predators may have been sufficiently incentivized to differentiate between the different phenotypes, and this is an important area for future work.

While our observed attack rates were lower than expected based on previous work conducted in the tropics (Finkbeiner et al., 2018; Seymoure & Aiello, 2015), they are comparable to several other experiments that used artificial models to assess avian predation (Kristiansen et al., 2018; Palmer et al., 2018; Robinson et al., 2025). We consider that these relatively low attack rates reflect our conservative predation scoring criteria and the use of a negative stimulus in the absence of a food reward. We also observed substantial variation in predator activity among field sites; for example, all sites in the high-density, no Bitrex treatment experienced at least one predation event, whereas predator activity was detected at only 19 of 25 field sites in the low-density, no Bitrex treatment. Although filtering our data to remove sites that experienced no predation and including site as a random effect in our model helps account for this heterogeneity, some uncertainty is inherent to field-based predation experiments. Our results highlight the potential for absolute density of mimetic communities to impact protection for imperfect Batesian mimics and provide a critical foundation for future work to leverage controlled experiments that could further disentangle the mechanisms underlying absolute density effects on predator foraging decisions.

Overall, our results align with previous work demonstrating that increased unpalatability of defended models enhances protection for aposematic prey and their imperfect mimics, and further suggest that, in complex natural environments with abundant alternative prey, the degree of mimetic perfection needed to benefit from enhanced warning signals may be broader than previously appreciated. Our findings also indicate that absolute density effects can strengthen warning signals in nature, driving stronger predator avoidance responses through mechanisms analogous to those associated with variation in model unpalatability and model–mimic relative frequencies (Endler & Mappes, 2004; Pfennig et al., 2001). Considered within the broader mimicry literature, this work underscores the importance of distinguishing between relative frequency and absolute density when testing hypotheses about relaxed selection in the context of mimetic prey. However, more work is needed to clarify how factors such as alternative prey availability, predator hunger, and the nutritional value of unpalatable prey modulate the effects of absolute prey density on protection for imperfect Batesian mimics.

## DATA AVAILABILITY

Data from the field predation experiments and code files for all analyses can be found at https://github.com/butterfliesrcool/Absolute_Density_Project.

## FUNDING

This work was funded by the National Science Foundation, Grant Number 2021181

## CONFLICT OF INTEREST

We declare no conflict of interests

## ACKNOWLEDGEMENTS

We would like to thank the Organization for Tropical Studies for allowing us to complete our experiments at La Selva Biological Station (Permit Number: SINAC-ACC-PI-re-023-2022) and Miguel Salazar and Jordan Smith for their assistance with fieldwork.

## Notes

https://doi.org/10.5061/dryad.sxksn03jk

https://github.com/butterfliesrcool/Absolute_Density_Project

### Competing Interest Statement

The authors have declared no competing interest.

### Summary of Updates

Author order and affiliations were updated

