## Supplemental Tables for "Increasing absolute prey community density protects aposematic models and their imperfect Batesian mimics: Evidence from Neotropical *Adelpha* butterflies"

**Table S1**. **(A)** Four mixed effects Cox proportional hazards models were fitted using backwards model selection to assess relative effects of species, absolute community density, and model unpalatably on model fit. All models included a random effect of site. **(B)** Models were then compared using likelihood ratio tests and AIC scores. We did not find any significant differences in model fit, so Model 3 was used in the final analysis to align with parsimony.

***A) Summary of fitted models***

Model response: Surv(experimental day, attacks)
**Model 1:** ~species * absolute density * model unpalatability + (1 | site)
**Model 2:** ~species + absolute density * model unpalatability + (1 | site)
**Model 3:** ~species + absolute density + model unpalatability + (1 | site)
**Model 4:** ~species * absolute density + model unpalatability + (1 | site)

***B) Results of model comparison***

| | **loglik** | **$X^2$** | **df** | **p-value** |
|---|---|---|---|---|
| Model 1 | -3071.6 | | | |
| Model 2 | -3074.0 | 4.7057 | 6 | 0.5821 |
| **Model 3** | **-3075.2** | **2.3788** | **1** | **0.1230** |
| Model 4 | -3074.3 | 1.7452 | 2 | 0.4179 |

**Table S2**. **(A)** Four generalized linear mixed models were fitted using the binomial distribution to assess significant predictors of attack rate on butterfly facsimiles. All models included a random effect of site. **(B)** Models were then compared using likelihood ratio tests and AIC scores. Models did not differ significantly in fit, so the simplest model (Model 3) was used in the final analysis to align with parsimony

***A) Summary of fitted models***

**Model 3:** attacks ~ species + model unpalatability + absolute density + (1 | site)
**Model 2:** attacks ~ species + model unpalatability * absolute density + (1 | site)
**Model 1:** attacks ~ species * model unpalatability * absolute density + (1 | site)

***B) Results of model selection***

| | **npar** | **AIC** | **BIC** | **loglik** | **-2*long(L)** | **$X^2$** | **DF** | **p value** |
|---|---|---|---|---|---|---|---|---|
| **Model 3** | **6** | **3485.2** | **3535.4** | **-1736.6** | **3473.2** | | | |
| Model 2 | 7 | 3484.8 | 3543.4 | -1735.4 | 3470.8 | 2.3657 | 1 | 0.1240 |
| Model 1 | 13 | 3492.1 | 3600.9 | -1733.0 | 3466.1 | 4.7467 | 6 | 0.5767 |
